# A juvenile climbing exercise establishes a muscle memory boosting the effects of exercise in adult rats

**DOI:** 10.1101/2021.06.02.446793

**Authors:** Einar Eftestøl, Eisuke Ochi, Inga Solgård Juvkam, Kristian Gundersen

**Author notes:** Corresponding author Visiting address: Department of Biosciences, University of Oslo, Kristine Bonnevies hus, Blindernveien 31, 0371 OSLO, Norway.

## Abstract

One of the ideas stemming from the discovery of a cellular memory in muscle cells has been that an early exercise period could induce a long-term muscle memory, boosting the effects of exercise later in life. In general muscles are more plastic in younger animals, so we devised a 5-week climbing exercise scheme with food reward administered to juvenile rats (post-natal week 4-9). The juvenile exercise increased fiber cross-sectional area (fCSA), and boosted nuclear accretion. Subsequently the animals were subjected to 10 weeks of detraining (week 9-19, standard caging). During this period fCSA became similar in the animals that had been climbing compared to Naive controls, but the elevated number of myonuclei induced by the climbing were maintained. When the Naive rats were subjected to two weeks of adult exercise (week 19-21) there was little effect on fCSA, while the previously trained rats displayed an increase of 19%. Similarly, when the rats were subjected to unilateral surgical overload in lieu of the adult climbing exercise, the increase in fCSA was 20% (juvenile climbing group) and 11% (Naive rats) compared to the contralateral leg. This demonstrated that juvenile exercise can establish a muscle memory. The juvenile climbing exercise with food reward led to leaner animals with lower body weight. These differences were to some extent maintained throughout the adult detraining period in spite of all animals being fed ad libitum, indicating a form of body weight memory.

## Introduction

Muscle memory related to strength training is referring to the phenomenon that if you were once strong, you could, even after a period of inactivity and loss of muscle mass, more easily re-acquire muscle mass. The term has also been used synonymously to motor learning (Gundersen 2016), and while motor learning might be important, a memory in the muscle cells themselves was recently demonstrated (Bruusgaard, Johansen et al. 2010, Egner, Bruusgaard et al. 2013, Lee, Kim et al. 2018). This “true” muscle memory, as opposed to motor learning, has been related to the finding that nuclei incorporated into muscle fibers during developmental and exercise-induced fiber growth are permanent (Bruusgaard and Gundersen 2008, Eftestøl, Psilander et al. 2020). Also, recent data suggests that the muscle fiber size is limited by the DNA content, i.e., the number of nuclei in these syncytial cells (Cramer, Prasad et al. 2020, Hansson, Eftestøl et al. 2020). This might reflect a general principle in cells (Gregory 2001, Gillooly, Hein et al. 2015).

A cellular memory related to DNA content could be very long lasting since nuclei extracted from human muscle tissue have a half-life of 15 years (Spalding, Bhardwaj et al. 2005), and since this includes a high proportion of stroma cells with high turnover, the myonuclei proper might be more stable or even permanent (for discussion see Gundersen 2016, Eftestøl, Psilander et al. 2020).

The mechanisms related to satellite cells providing new myonuclei is impaired during the lifespan (Schultz and Lipton 1982, Chen, Datzkiw et al. 2020). This has led to the idea that you should exercise when young and benefit later (Gundersen, Bruusgaard et al. 2018). Such an insight would have profound influence on public health advices to a population where sarcopenia is an increasing problem (Dutta and Hadley 1995, Hughes and Schiaffino 1999).

We therefore wanted to investigate whether an exercise period in young rats induced a muscle memory that boosted effects of retraining in adult rats even after a prolonged intervening detraining period. The initial exercise period started at 4 weeks of age, at a time of active growth-related myonuclear accretion, and largely before the number of myonuclei per mm fiber reaches a plateau (Enesco and Puddy 1964, Cheek, Holt et al. 1971, Schultz 1996, White, Bierinx et al. 2010). At the age of four weeks the body weight is <20% (Sengupta 2013) and bone length <50% (Reichling and German 2000) compared to fully grown rats. The surge of testosterone connected to puberty occurs at week 7-9 (Ojeda and Urbanski 1994, Lewis, Barnett et al. 2002, Picut, Remick et al. 2015).

We here provide data suggesting that climbing exercise in juvenile rats (4-9 weeks postnatally) boosts effects of subsequent exercise in adult rats.

## Methods

### Animals and ethical considerations

Male Sprague Dawley rats with age 4 weeks at experiment start were used. Rats were kept at the animal facility at the Department of Biosciences, University of Oslo, housed with a 12 h light/dark cycle. Rats were euthanized by cervical dislocation while under deep isoflurane anesthesia. All animal experiments were conducted in accordance with the Norwegian Animal Welfare Act of 20th December 1974, and all experiments were approved by the Norwegian Animal Research Committee before initiation. The Norwegian Animal Research Authority provided governance to ensure that facilities and experiments were in accordance with the Act, National Regulations of January 15th, 1996, and the European Convention for the Protection of Vertebrate Animals Used for Experimental and Other Scientific Purposes of March 18th, 1986.

### Rat exercise protocol

Rats were randomly assigned to either a “Memory” or a “Naïve” group at experiment start, where Memory rats were housed in a custom-made climbing cage for an initial 5-week period (1. training), where the inner walls and ceiling were covered with chicken wire. The food pellets were placed in a chicken wire container in the middle of the ceiling, so rats needed to climb and cling to the ceiling in order to feed. Water was accessible at all times without climbing. Naive rats were housed in standard cages for this initial training period with ad libitum access to food and water. Per-cage food intake and individual body weights were measured weekly during 1. training and detraining, and then on week days during retraining by climbing. Three separate experimental series were performed in this paper: Series 1) 4-week-old rats were housed either in climbing cages (Memory) or in standard cages (Naive) for 5 weeks (1. training period) before termination. Series 2) A second batch of rats followed the same initial training setup as in Series 1, followed by a detraining period in standard housing for 10 weeks (detraining period), and a consecutive retraining period (RT) in climbing cages for 2 weeks (Memory). Age-matched controls were housed in standard cages up until retraining start, and then housed in climbing-cages for 2 weeks (Naive), simultaneously with the RT period for Memory rats. One group of Memory and Naive rats were terminated at the end of the detraining period, and another group were assigned to the climbing cages, either for the first time (Naive) or for the second time (Memory). No rats were terminated after the 1. training period for this series. Memory and Naive groups were at the end of the detraining period assigned cage-wise to best match average body weight between groups within the detraining and retraining groups, respectively. Series 3) A third batch of rats performed a 1. training and detraining period as in series 2, followed by a two-week period with unilateral overload of the soleus muscle in the right leg by removing 1/3 of the distal part of the gastrocnemius. This in order to infer whether overload would lead to an increased hypertrophic response relative to its contralateral leg with identical history in Memory rats compared to in Naive rats. A timeline illustration of the experimental setup relative to developmental phases is shown in Fig. 1.

**Fig.1:**
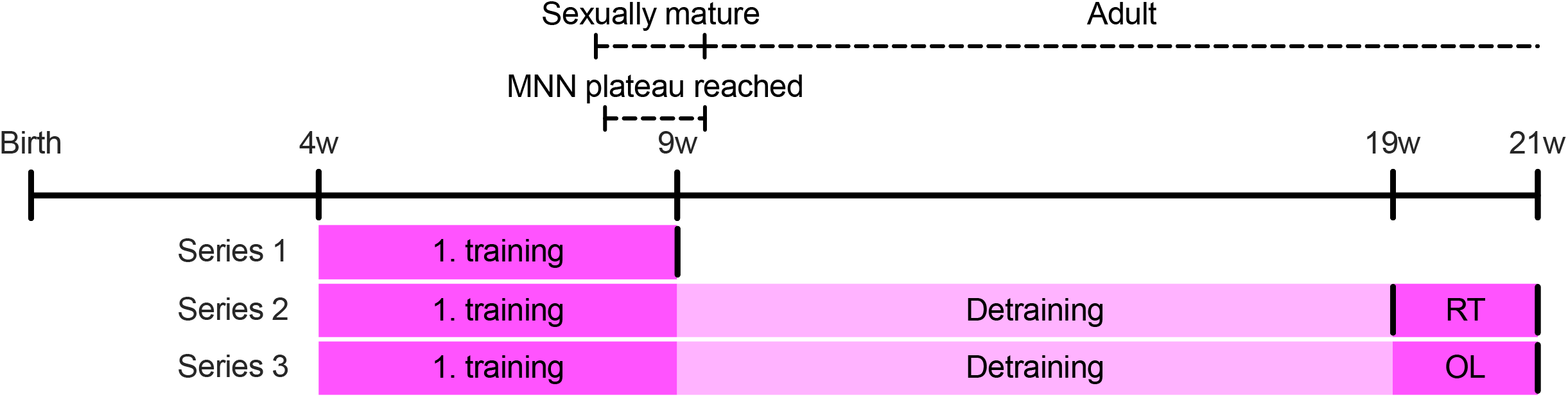
Rat experimental setup in light of relevant developmental checkpoints. Rats were introduced to a climbing cage (1. training) 4 weeks (w) after birth, followed by a 10w Detraining period. Retraining in climbing cage (RT), or by unilateral overload (OL) of the soleus muscle, was started at age 19w. Results are based on three separate experimental series, vertical lines representing termination time-points of rats from the respective series that performed the initial climbing period, referred to as the “Memory” group throughout. Age-matched “Naive” controls not performing the first climbing period, thus being “Naive” when introduced to the climbing cage or OL at age 19w, were included for all series. Results from the tibialis anterior (TA) muscle for Series 1 and 2, and from the soleus for Series 3 is presented throughout. Dashed lines with text showing time-period for when male rats reach sexual maturity and myonuclear number (MNN) reaches a plateau phase (see references in main text).

At timepoints after 1. training, detraining and retraining, rats were sedated with 2% isoflurane, weighed, and the respective muscles were isolated (tibialis anterior (TA) for series 1 and 2, soleus for series 3), blotted for excessive fluid, weighed, covered in Tissue-Tek O.C.T. compound (Sakura), frozen in isopentane cooled in liquid nitrogen and then stored at -80 °C until further analysis. Rats were then terminated, and for experimental series 2, intra-abdominal fat content was also excised and weighed.

### Immunohistochemistry

Muscles embedded in OCT were equilibrated in a freezing microtome (Leica CM1950) and sectioned at 10 μm. Sections from Memory and Naive muscles within each time-point and experimental series were always placed side-by-side on slides to obtain standardized staining and imaging conditions between groups. Slides were stored in -80 °C until further processing.

A weakness with previous studies on muscle cross sections has been the criteria used to determine myonuclei from other cell nuclei. By basing the determination of true myonuclei purely on their location in relation to the boundary of the muscle fiber, it may lead to false positives or negatives (Bruusgaard et al., 2012; Bruusgaard et al., 2010). To overcome this problem, herein we used an antibody against pericentriolar material 1 (PCM1) that specifically labels myonuclei (Winje, Bengtsen et al. 2018). Sections were retrieved from -80 °C and equilibrated for 30 min at room temperature. Prior to staining, sections were pre-incubated with 2% bovine serum albumin (BSA) in PBS pH 7.4 for 30 min. Sections were stained with a rabbit primary antibody against PCM1 (1:1000, HPA023370, Sigma-Aldrich) in staining solution (5% BSA in PBS pH 7.4, 0.2% Igepal CA-630) overnight at 4 °C. Next day the sections were washed 3 × 10 minutes with PBS and stained with an anti-rabbit secondary antibody (1:1000, AB150077, Abcam, Alexa 488) in 2% BSA in PBS for 1 hour. Sections were washed 3 × 10 minutes in PBS then stained with a mouse primary antibody against dystrophin (1:20, MANDYS8, 8H11, DSHB) in staining solution as described above. The sections were again washed 3× 10 minutes with PBS and stained with an anti-mouse secondary antibody (1:500, A-11005, Thermo Fisher Scientific, Alexa 594) in 2% BSA in PBS for 1 hour. Sections were washed 3 × 10 minutes with PBS and mounted with DAPI Fluoromount-G (Southern Biotech, cat. 0100-20).

### Imaging and image analysis

Immunostained cryosections were visualized on a FluoView FV 1000 Olympus inverted confocal microscope, with a resolution of 1024×1024 pixels (317,44 × 317,44 μm) and dwell time of 2 μs/pixel using a 40x PlanApo oil immersion objective (NA 1.3). Laser with excitation wavelength peak at 408 nm was used to excite DAPI to visualize all nuclei. A 488 nm laser was used to visualize myonuclei (PCM1). A 594 nm laser was used to visualize the border of the muscle fibers (dystrophin). All system settings were standardized between groups. Image analysis were performed using Adobe Photoshop CS6 (Adobe Systems, USA). 6-15 images were acquired from each muscle, and a total of 93-190 fibers per muscle were analyzed. Fibers with their entire dystrophin ring inside of the image frame were counted, excluding fibers with central nuclei or inconsistent dystrophin ring due to focal plane differences. Nuclei with both PCM1 and DAPI positive staining were counted as myonuclei. Fiber cross-sectional area (fCSA) was analyzed based on intensity differences between the inside of the fiber and the inner rim of the dystrophin staining, using the magic want tool and/or quick selection tool, and manually adjusted when incorrect.

### Statistics

Statistical analyses were performed in Prism 8 for macOS (Version 8.4.3, GraphPad Software, LLC). A nested t test for each time-point was performed between Naive and Memory rats for analysis of fCSA (Fig. 2B) and myonuclear number (Fig.3A) from series 1 and 2. A Wilcoxon matched-pairs signed rank test was performed for within-rat comparisons of muscle wet weight, fCSA (muscle average values) (Fig. 2D) and myonuclear number (Fig. 3C) between the overloaded and contralateral control leg. Myonuclear domain (MND) was analyzed with a One-way ANOVA with Tukey’s multiple comparisons test with a single pooled variance (Fig. 4A). A nonlinear regression analysis fitting a straight line using least squares regression, comparing slopes with the Extra sum-of-squares F test, was used for the correlation analysis between MNN and fCSA (Fig. 4B). Terminal body weight, intraperitoneal fat content and muscle wet weight (left and right leg average) were analyzed with a Mann-Whitney test for each parameter between Naive and Memory rats in experimental series 1 (Fig. 5A). Timeline data (body weight and food intake) were analyzed with a Mixed effect model (REML) with Sidak’s multiple comparisons post hoc test between Naive and Memory rats at each time-point (Fig.5B and Suppl. Fig.1). Result section values are given as muscle average grand mean +/-SD. Values and number of biological replicates represented in figures are specified in figure legends.

**Fig.2:**
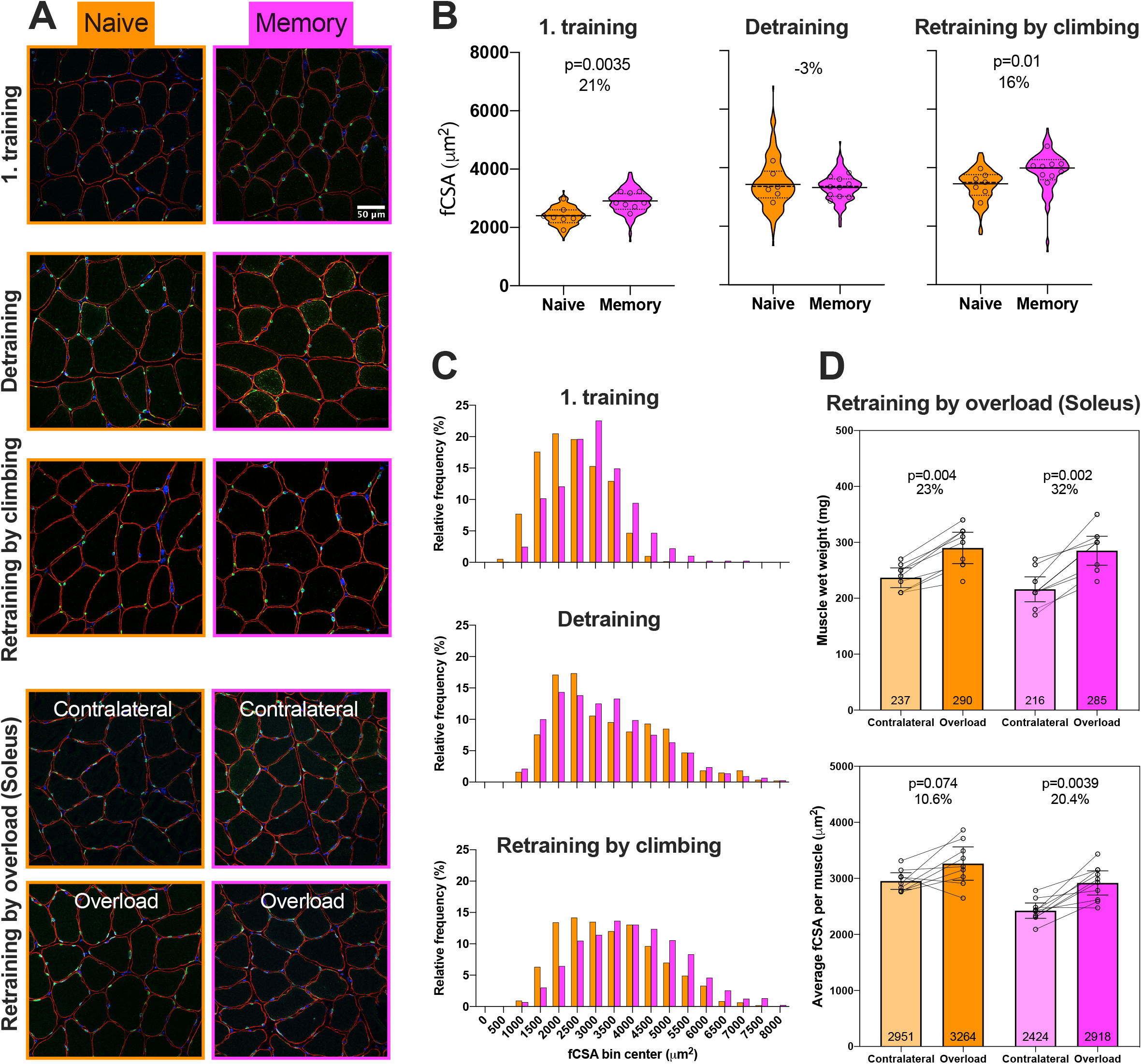
Training when young facilitate hypertrophy when re-introduced to the same training or synergist-ablation induced overload when adult. (A) Example images of TA or soleus cross sections showing fiber boundaries (dystrophin, red), myonuclei (PCM1, green) and all nuclei (DAPI, blue). Equal scale for all images. (B) Violin plots of pooled individual fiber cross-sectional area (fCSA) values including median (dashed lines) and quartiles (dotted lines) superimposed with per muscle mean values (circles) including the muscle grand mean (long full lines). (C) Frequency distributions of individual fiber fCSA. (D) Group mean (bars, number inside bars, 95% CI) muscle wet weight (top) or fCSA (bottom), per muscle mean values (circles) and within-rat contralateral comparisons (lines). Number of fibers/muscles (n/N) in Naive;Memory group: 1. training) 1307/8;1259/8, Detraining) 876/6;1526/10, Retraining by climbing) 1058/7;1439/9, Retraining by overload) Contralateral: 734/9;1278/10, Overload: 2203/9;2638/10.

**Fig.3:**
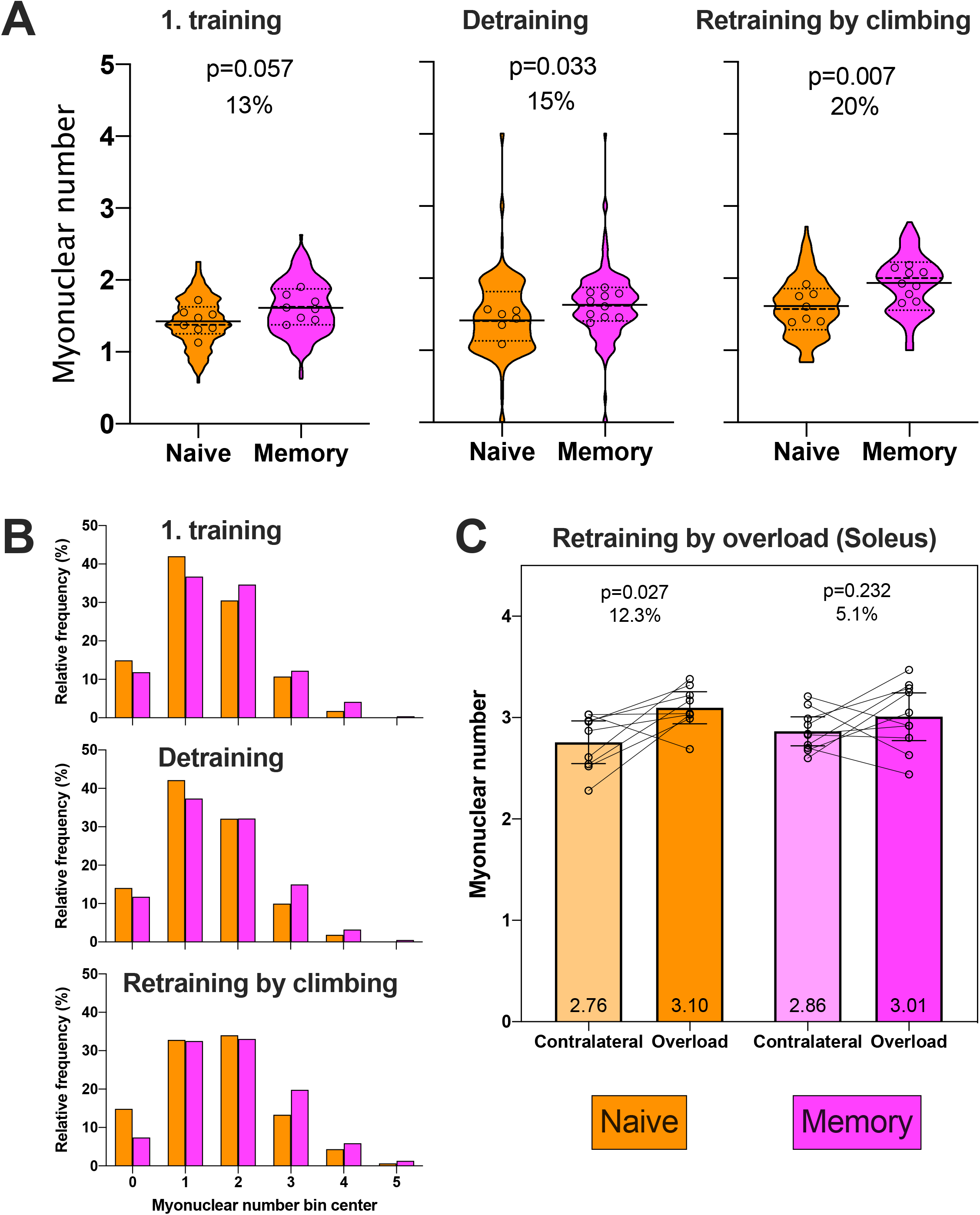
Training when young facilitate myonuclear accretion, and myonuclei are retained into adulthood after a period of detraining. (A) Violin plots of pooled individual fiber myonuclear number from muscle cross sections including median (dashed lines) and quartiles (dotted lines) superimposed with individual muscle mean values (circles) including the muscle grand mean (long full lines). (B) Frequency distributions of myonuclear number per fiber cross section. (C) Group mean myonuclear number per muscle (bars, number inside bars, 95% CI), per muscle mean values (circles) and within-rat contralateral comparisons (lines). Number of fibers/muscles (n/N) in Naive;Memory group: 1. training) 1307/8;1259/8, Detraining) 876/6;1526/10, Retraining by climbing) 1058/7;1439/9, Retraining by overload) Contralateral: 734/9;1278/10, Overload: 2203/9;2638/10.

**Fig.4:**
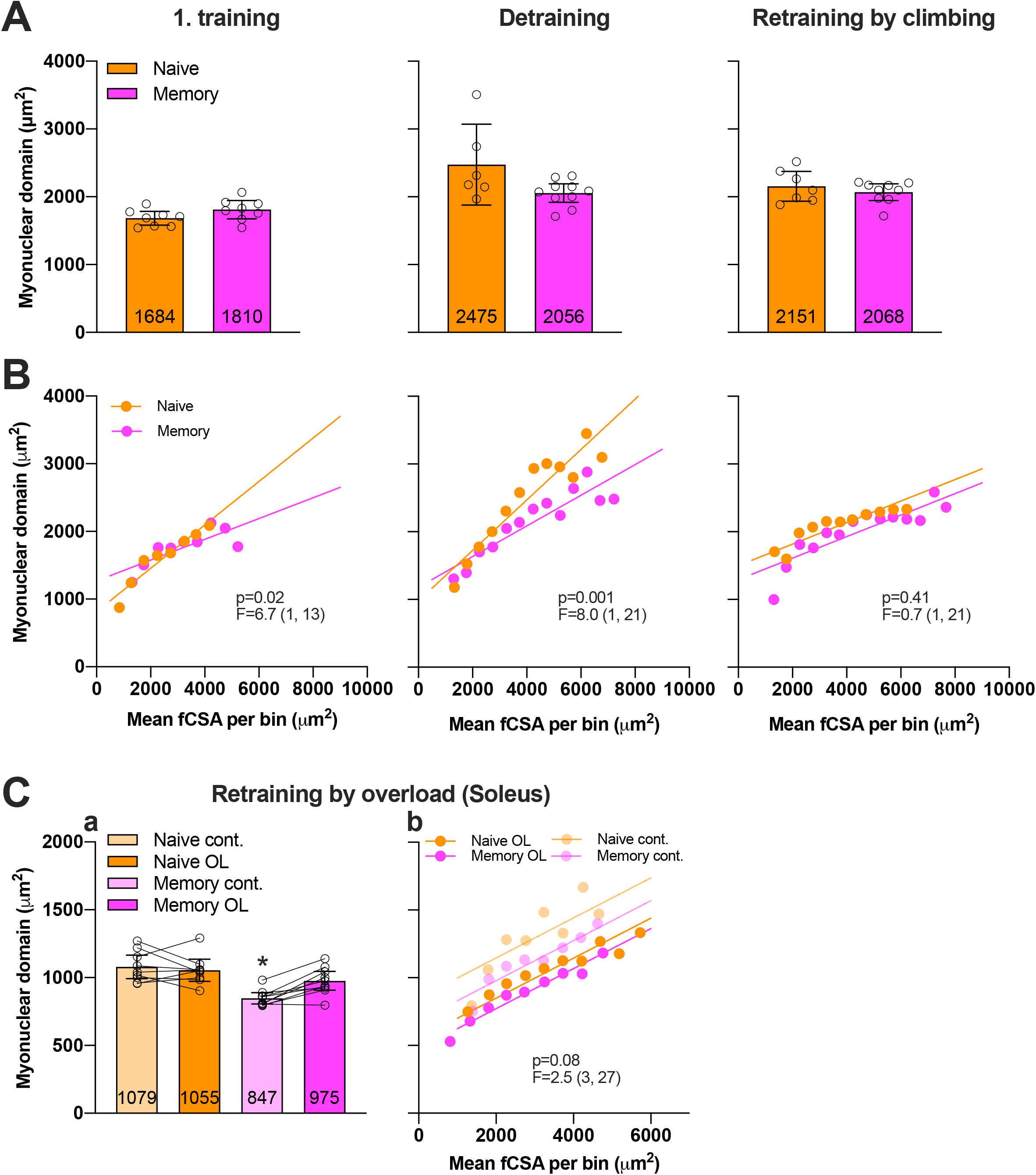
Myonuclear domain size is dictated by the number of myonuclei during hypertrophy and atrophy. (A, Ca) Myonuclear domain size shown as group mean (bars), muscle mean (circles), 95% CI and comparisons between overloaded (OL) and non-overloaded contralateral control leg (cont.)(before-after lines)(Ca). (B, Cb) Correlation between mean fiber CSA (fCSA) within bins with size of 500µm^2^ vs corresponding mean myonuclear domain size from the same fibers within each fCSA bin (circles) and regression lines. * denotes different from all other groups (p≤0.025). p- and F (DFn, DFd)-values denote slope comparisons from a nonlinear regression analysis. Number of fibers/muscles (n/N) in Naive;Memory group: 1. training) 1302/8;1236/8, Detraining) 858/6;1521/10, Retraining by climbing) 1046/7;1284/9, Retraining by overload) Contralateral: 734/9;1278/10, Overload: 2203/9;2638/10.

**Fig.5:**
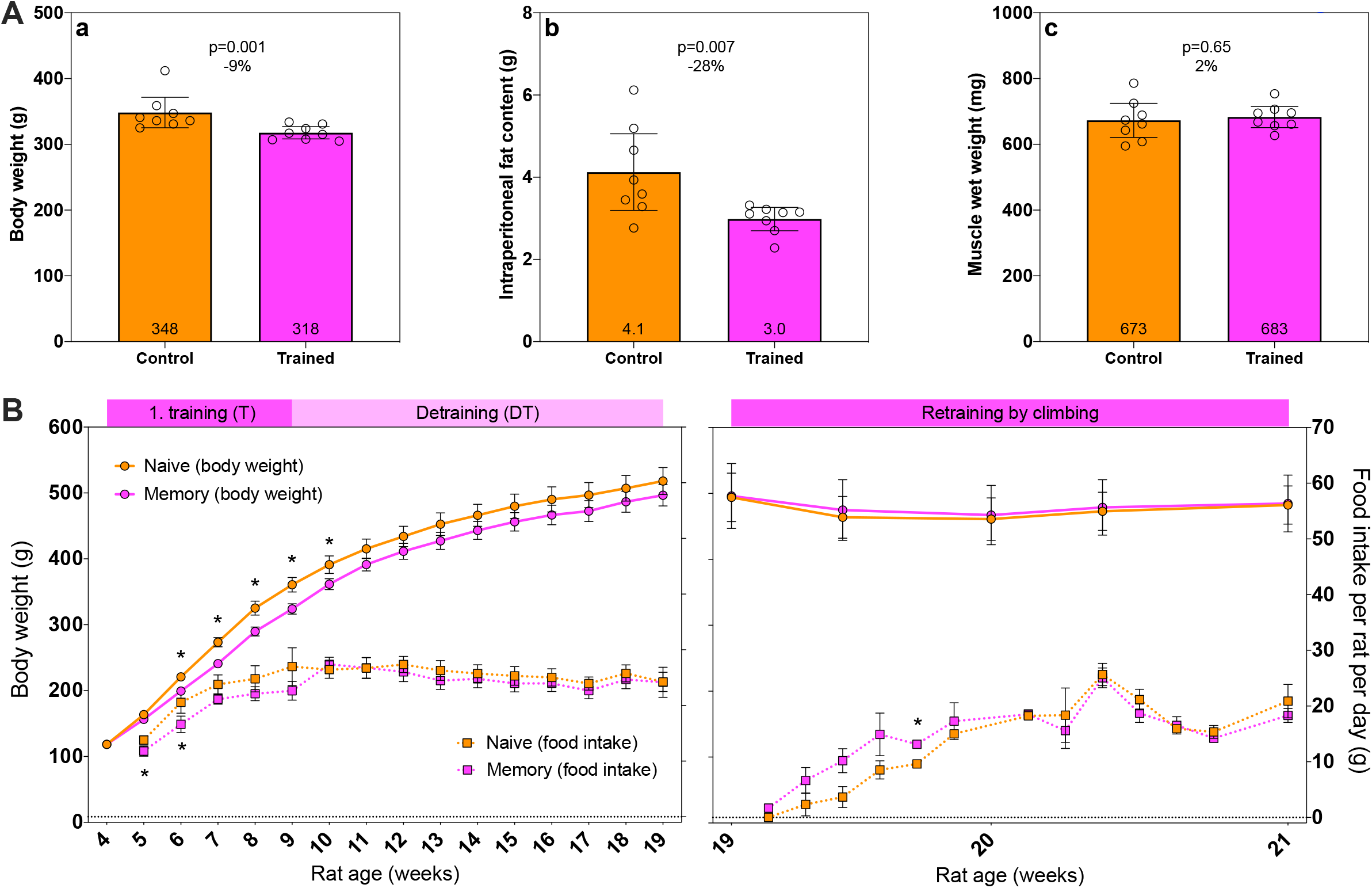
Effect of training, detraining and retraining on body weight, food intake, fat content and muscle weight. (A) Body weight (a), intraperitoneal fat content (b) and muscle wet weight (c) showing group mean (bars, numbers inside bars, 95% CI) and individual rat values (circles) after the initial 5-week climbing period from experimental series 1. (B) Timeline changes in body weight (circles connected with lines) and food intake (squares connected with dotted lines) during the 1. training, detraining (mean, 95% CI) and retraining period (mean, SD) from experimental Series 2. * denote significant difference between Naive and Memory rats at a given time-point. Body weight N (rats)/food intake N (cages): 1. training and detraining; Memory N=30/9, Naive N=20/6. Retraining; Memory N=10/3, Naive N=7/2.

## Results

The experiments involved a juvenile climbing exercise period (1. training) (postnatal week 4-9), a 10-week detraining period under standard caging conditions (week 9-19), and a subsequent 2-week adult retraining period either by climbing or by synergist ablation. The experiments were performed as three separate experimental series as illustrated in Fig. 1.

### Climbing exercise in young rats facilitated a hypertrophy-response to adult exercise

In the first series 4w old rats were subjected to 5 weeks of climbing exercise (“Memory” group), where the tibialis anterior (TA) was analyzed at week 9. The training led to an average increase in fiber cross-sectional area (fCSA) of 21% (p=0,004) compared to controls subjected to standard caging (“Naive” group; (Fig. 2B)). After a subsequent detraining period (Series 2) where both groups were caged under standard conditions for 10 weeks and ending at 19 weeks age, the Naive and Memory rats reached similar fCSA (Fig. 2B-C). Retraining by climbing for 2 weeks at this time had little effect on Naive rats, while the Memory rats displayed an increase in average fCSA of 19% which was 16% higher than in the Naive group (p=0.01; Fig. 2B). These effects was also reflected in the size distribution at the individual fiber level by a general shift to larger fibers in the Memory group after 1. training and retraining compared to the Naive group (Fig. 2C). This clearly demonstrated a memory effect such that the juvenile climbing exercise enhanced the effects of the adult training with respect to hypertrophy.

A third experimental series (Series 3) were similarly subjected to juvenile climbing and subsequent detraining. But for the retraining, the gastrocnemius muscle was partly ablated in one leg in both Naive and Memory rats in order to overload the soleus muscle which was used in the subsequent analysis. In this series, the Memory group had in general smaller fCSA than the Naive group (Fig. 2D), possibly due to lower body weights (discussed below), carried by the postural soleus. When the overloaded soleus in the Naive rats were compared to the contralateral leg the average fCSA was 11% higher, while in the Memory rats it was 20% higher (Fig. 2D). Muscle wet weights essentially reflected the fCSA changes (Fig. 2D). This further supported the notion of a muscle memory where juvenile exercise boosted the effects of the adult overload.

### Climbing exercise in young rats led to a permanent increase in the number of myonuclei

The juvenile climbing exercise led to an increase in the number of myonuclei by 13% compared to sedentary controls (p=0.057; Fig. 3A). Although this was only borderline to statistically significant, after the detraining period the difference between the climbing and sedentary group was essentially maintained and was statistically significant (15% difference, p=0.033) (Fig. 3A). Retraining by climbing for 2 weeks increased the number of myonuclei by 19% in Memory- and 14% in Naive rats, resulting in the Memory rats having on average 20% more nuclei compared to the Naive group (p=0.007; Fig. 3A). These effects was also reflected in the frequency distribution at the individual fiber level by a general shift to increased number of myonuclei in the Memory group compared to the Naive group (Fig. 3B). These observations support the idea that the permanent increase in the number of nuclei could act as a functional substrate for the muscle memory and explain the difference in the training responses between the Naive and Memory rats.

Myonuclear number did not increase in either Naive or Memory rats during the detraining period (week 9-19), indicating that the growth related myonuclear accretion reached its plateau phase before this period.

When the soleus muscle was subjected to unilateral overload, as a substitute for the retraining by climbing, myonuclear number was increased by 12% in the overloaded leg in Naive rats, and by 5% (not significant) in the Memory rats when compared to their respective contralateral legs (Fig.3C).

### Establishment of a muscle memory did not alter myonuclear domain volume (MND)

The myonuclear domain (MND) was not altered by juvenile climb-exercise (Fig. 4A), suggesting that the increase in fCSA was closely related to the number of nuclei. It should be noted however that for all experimental groups MND increased with fiber size (Fig. 4B-C). Thus, there was not an absolute MND per nucleus, but the MND was similar for fibers of the same size whether they had been exercised or not.

There was no change in the number of nuclei during detraining, but the increased fCSA led to larger MNDs. This was particularly prominent in the Naive group (47%), and less so in the Memory group (14%) (Fig. 4A). Plotting MND against fCSA, revealed that the difference in MND between the two experimental groups after detraining increased with increasing fiber size (Fig. 4B).

During retraining by climbing the memory group fCSA increased considerably and the two groups displayed similar MND and a similar slope between MND and fCSA (Fig. 4B). Similar observations were made after OL (Fig. 4Cii). Thus, the relationship between the number of nuclei and fiber size was similar for the Naive and Memory groups after the retraining by climbing or overload, and fibers of similar size had similar MNDs irrespective of their history.

### Juvenile training reduced food intake, body weight and fat content, and lower weight was preserved into adulthood

In the animals in series 1, the terminal body composition was determined after the juvenile training period and compared to sedentary controls. Training led to a 9% (p=0.001) lower body weight, 28% (p=0.007) lower intraperitoneal fat content, while muscle mass was similar in the two groups (Fig.5A).

When rats were followed longitudinally in Series 2 throughout the juvenile climbing period, they had a 16% lower total food intake compared to sedentary controls, and broken down on individual timepoints the intake was statistically different for the first two weeks (Fig. 5B). Note that food intake measurements were limited to per-cage averages. The body weight was lower in Memory rats and the difference was statistically significant after the 2. week of climbing through the first week of detraining (Fig.5B). At the end of the juvenile training period Memory rats had a 10% lower mean body weight compared to Naive rats. Only small differences in food intake were evident throughout the detraining period, and the difference in body weight between Memory and Naive rats obtained during the 4-9 weeks climbing period was gradually decreased throughout this period (Fig. 5B). At the end of detraining, average body weight was 4% lower in Memory rats compared to Naive rats.

A subset of the Naive and Memory rats from Series 2 were selected for similarity in body weight at the end of the detraining period in order to study rats of the same size during the adult training period (Fig. 5B). Neither group had significant changes in body weight during retraining (Fig. 5B). Both groups had a very low food intake in the beginning of the retraining period, but displayed a steep increase in food intake during the first week of exercise (Fig. 5B), in particular in the juvenile training group. Thereafter the food intake seemed to stabilize at a level somewhat lower than when food was freely available during detraining, and there were no significant differences between the two groups (Fig. 5B).

In Series 3 the climbing rats gained 464% and the sedentary controls 517% in body weight from experiment start to after overload, and had a steeper growth curve than in Series 2. No selection of body weight was done, and the difference in body weight was maintained throughout the detraining and overload retraining period (p<0.01 for all three time-points) (Suppl. Fig. 1).

## Discussion

We found that a juvenile climbing exercise led to 21 % increase in fCSA compared to sedentary controls, and even if the effect on fiber size was completely lost during subsequent detraining, the juvenile training boosted the effects of retraining on fCSA in the adult 10 weeks later. Juvenile training was accompanied by a permanent increase in the number of myonuclei that might be the functional substrate for the muscle memory (Bruusgaard, Johansen et al. 2010, Bruusgaard, Egner et al. 2012, Egner, Bruusgaard et al. 2013, Lee, Hong et al. 2016).

Theoretically, fiber size is related to the number of nuclei, and the ability of each nucleus to produce volume. The latter could be viewed as a flexibility in the MND, and was demonstrated by the finding that MND increases with fiber size (Fig. 4) (Hansson, Eftestøl et al. 2020). As we have suggested previously such flexibility might be related to either the number of nuclei limiting the maximum fiber size (Hansson, Eftestøl et al. 2020), or the presence of some form of cooperativity between nuclei increasing their ability to produce volume (Cramer, Prasad et al. 2020).

However, the relationship between the MND and fiber size was the same after the 1. training and retraining, and when comparing the Memory groups to the Naive controls for a given fiber size. Thus, our data fit a model where the number of myonuclei dictates fiber size, and where the memory effect is related solely to the number of nuclei, and not to the ability of each nucleus to produce volume.

The experiments with overload after synergist ablation in lieu of a 2. climbing exercise showed essentially the same result. This is important, because the better effect of a 2. climbing exercise might be influenced by motor learning rather than a cell autonomous muscle cell memory. This seems less likely for the memory effect revealed by overload.

The juvenile exercise with food reward, reduced food intake, body weight and body fat content. Interestingly the difference in weight persisted throughout the detraining period in spite of all animals being offered food ad libitum. This suggests the presence of a “body weight memory”.

In a previous study on muscle memory in somewhat older female rats (week 8-16) Lee et al. (Lee, Kim et al. 2018) used a weight loaded-ladder for exercise and observed a modest increase in fCSA of 9% (compared to our 21%), but their increase in the number of myonuclei was much larger, 23% compared to our 13%. Similar to our findings, detraining led to a complete loss of the exercise effect on fCSA, with no loss of myonuclei. After eight weeks of retraining (age week 36-44) their study showed a fCSA increase similar to ours, 15%.

The differences in sex, training duration and protocol makes it hard to compare the two studies, but it is noteworthy that the relative increase in the number of nuclei induced by the first training session was larger in the older animals.

A priori we had expected juvenile exercise to be particularly potent increasing nuclear accretion, as the potential for myonuclear accretion in developing rats is high (Moss and Leblond 1970, Moss and Leblond 1971, Schiaffino, Bormioli et al. 1976), and the proliferative potential of satellite cells is reduced with age (Schultz and Lipton 1982). Our data indicated however, that while juvenile exercise was efficient in establishing a memory for fCSA, it was not particularly efficient in boosting nuclear accretion. We speculate that when exercising at a time when myonuclear accretion is very high due to normal growth even in sedentary animals, the capacity for further accretion induced by exercise might be limited. Our results confirmed, however, that there is a strong correlation between myonuclear number and fiber size, and that an increased myonuclear number is still maintained upon detraining even in developing adolescent rats where basal nuclear insertion is ongoing.

Our results do not exclude that additional mechanisms other than an elevated number of nuclei could explain the induction of a muscle memory such as long-term changes in motor behaviour (Psilander, Eftestol et al. 2019) and epigenetic changes (Seaborne, Strauss et al. 2018).

Juvenile strength training in humans are deemed safe (Dahab and McCambridge 2009), and has effects similar to those in adults on voluntary strength (Ramsay, Blimkie et al. 1990). It has however been unclear to what extent such exercise is affecting the muscle cells or if it is largely an effect of motor learning (Ramsay, Blimkie et al. 1990). Thus, most studies have reported little effect on anthropometrical muscle cross sectional area (Ramsay, Blimkie et al. 1990). In contrast, a MR study on twins suggested that static training increased lean muscle CSA (Mersch and Stoboy 1989). Moreover, evoked twitch-strength can be enhanced by juvenile exercise suggesting some form of muscle cell adaptation (Ramsay, Blimkie et al. 1990). Irrespective of mechanism, a longitudinal study comparing children with and without a juvenile strength exercise would reveal if the imprinted muscle memory that we have uncovered in juvenile rats could provide long-term effects on trainability and health in adults, and thus, if such exercise should be emphasized in the school curriculum.

## FIGURE LEGENDS

**Suppl. Fig.1:**
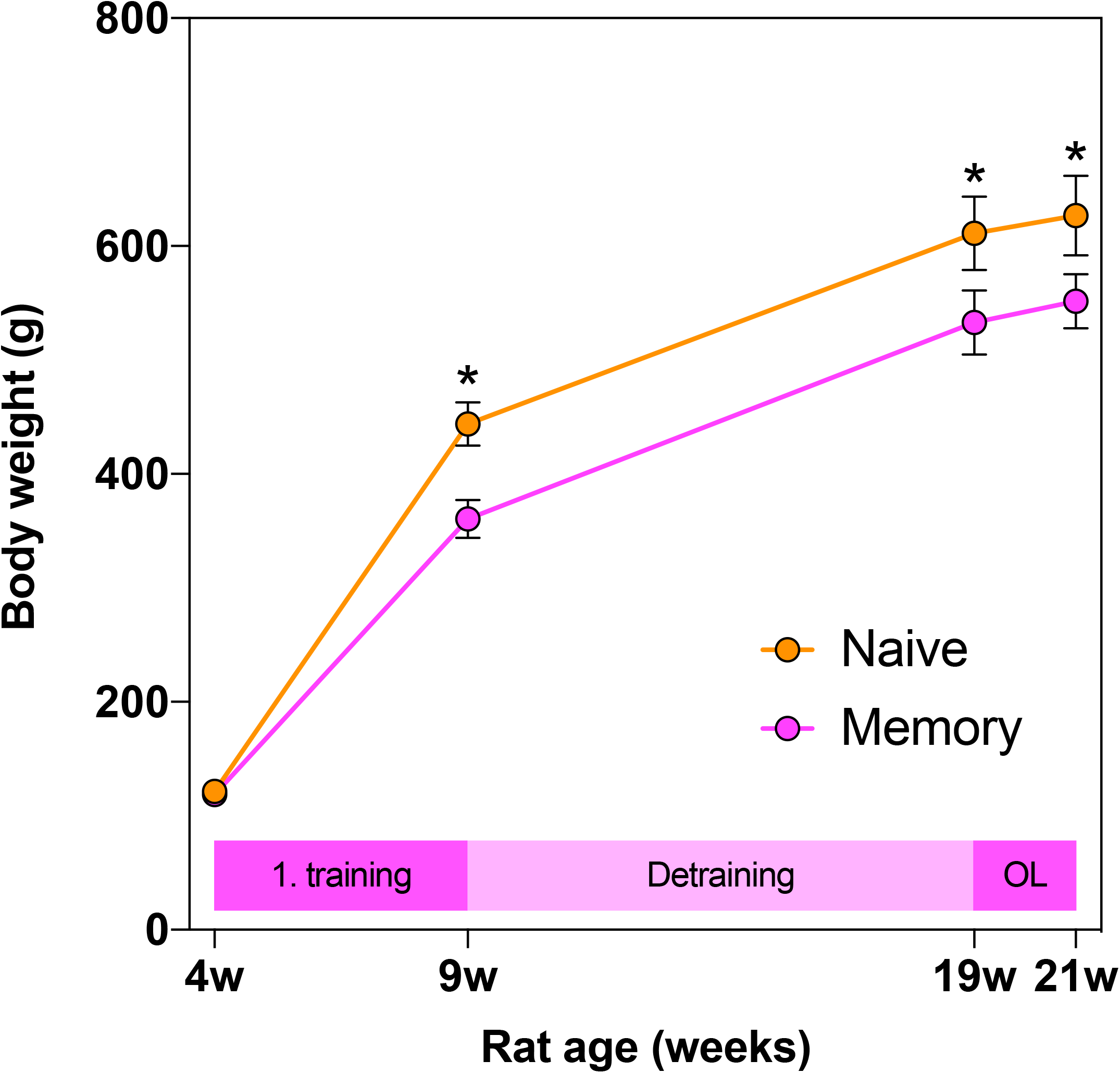
Effect of training, detraining and unilateral overload on body weight. Timeline changes in body weight (circles connected with lines, group mean, 95% CI) during the 1. training, detraining and unilateral overload (OL) period from experimental Series 3. * denote significant difference between Naive and Memory rats at a given time-point. Naive N=9, Memory N=10.

**Suppl. Fig.2:**
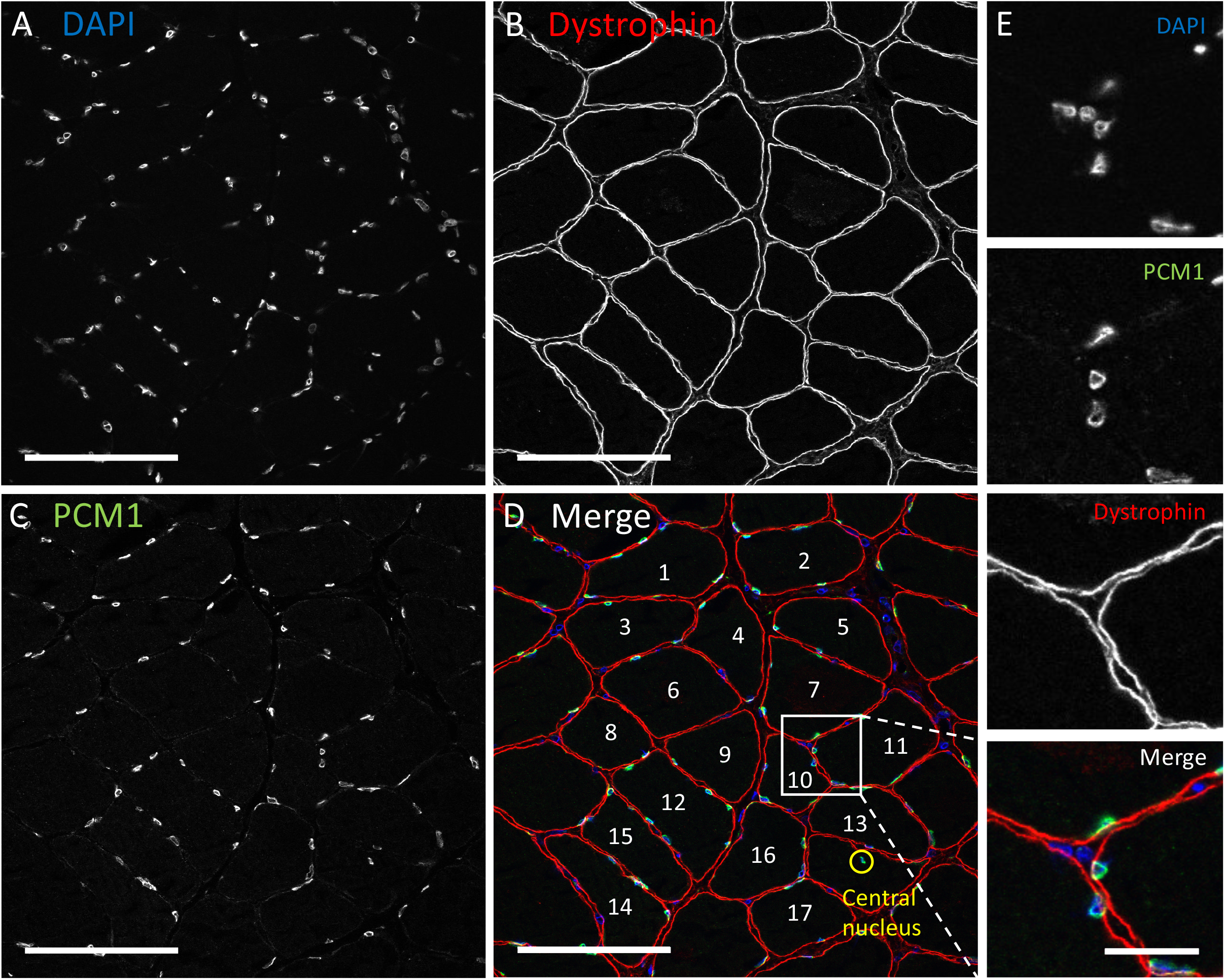
Immunohistochemically stained muscle cross section. Example of a muscle cryo cross-section (thickness 10 µm) used for analysis, showing (A) nuclei (DAPI), (B) muscle fibers (dystrophin), (C) myonuclei (PCM1) and (D) all three channels merged. The myonucleus marked with a yellow circle represents a central nucleus. Scale bar: 100 µm. (E) Enlarged view of A-D. Scale bar: 20 µm.

